# Molecular cloning and characterization of an AP2/ERF protein gene in cotton (*Gossypium hirsutum* L.)

**DOI:** 10.1101/2023.04.27.538633

**Authors:** Chaofeng Wu, Xuemei Ma, Shuyan Li

**Author notes:** Corresponding author Address:Anyang Institute of Technology Anyang Henan, 455000, China. Chaofeng Wu, Shuyan Li contributed equally to this work.

## Abstract

Upland cotton (*Gossypium hirsutum* L.) is one of the most economically important crops worldwide due to the significant source of natural fiber, feed, oil and biofuel products. Cottonseed can also serve as an excellent source of edible protein and oil. However,the presence of gossypol in pigment gland has limited it utilization In the past few decades, some progress has been made in the understanding molecular mechanism of the formation of the pigment gland. However, little is known about the specific mechanism of pigment gland formation in cotton. In this study, the cDNA sequence of a ethylene transcription factor gene, designated GhERF105a, was cloned from upland cotton CCRI12. Sequence alignment revealed that GhERF105a gene contained a typical AP2/ERF domain of 61 amino acids, and belonged to the ERF subgroup of the ERF supfamily. It was highly expressed in the leaves and stems of glanded plants but had substantially lower expression of the glandless plants. GhERF105a, localized to the nucleus, could bind to GCC-box and DRE. Some development, phytohormone and stress related cis-elements were enriched in the promoters of *GhERF105a/d*. Split ubiquitin assays in yeast and BiFC experiments showed extensive interactions between GhERF105a and Gh_A07G1044. In addition, *GhERF105a* was highly similar with *GhERF105d* in the gene length, molecular weight, protein molecule, gene structure and expression pattern. The overall results suggested that *GhERF105a* might participate in the pigment gland formation and stree-response processes.

## 1. Introduction

Upland cotton (*Gossypium hirsutum* L.) is not only the world’s leading textile fiber and oilseed crop, but also a crop that is of significant value for feed, foodstuff and biofuel products. In addition, cottonseed contains a relatively high-quality protein which can meet the annual protein requirement of half billion people (Sunilkumar et al. 2006; Gao et al. 2010). However, the use of cottonseed as protein source is limited due to the presence of gossypol stored in pigment glands that is toxic to human beings and monogastric animals (Bell and Stipanovic. 1977; Sang et al. 1980; Shandilya et al. 1982; Weinbauer et al. 1983; Heywood. 1988; Du et al. 2004; Stipanovic et al. 2006; Zhang et al. 2007). On the other hand, gossypol, as phytoalexin, can protect plants from pathogens, insects, and herbivores in certain species of cotton plants of the family Malvaceae (Bell and Stipanovic. 1977; Stipanovic et al. 1978a; Stipanovicet al. 1978b; Hedin et al. 1992; Wang et al. 2004; Townsend et al. 2005; Mao et al. 2007; Cai et al. 2010; Gao et al. 2013; Tian et al. 2018). In addition, it has of anti-tumor activity and contraceptive properties (Blackstaffe et al. 1977; Matlin 1994). Therefore, a better understanding of cotton pigment gland formation and gossypol biosynthesis regulation will greatly facilitate to develop cotton varieties with low - gossypol seeds and high - gossypol plants for researchers.

Research on the molecular genetic mechanisms of pigment gland began in the 1950s (McMichaelet al. 1954; McMichael et al. 1959; McMichael et al. 1960). Since then, gland formation is controlled by a combination of at least six independent loci such as *gl1*, *gl2*, *gl3*, *gl4*, *gl5* and *gl6*, the different combinations of dominant (*Gl*) and recessive (*gl*) alleles regulate gland formation in different organs (McMichael et al. 1960; Lusas and Jividen. 1987; Lee 1965; Gutierrez et al. 1972; McCarty et al. 1996; Scheffler et al. 2008; Scheffleret al.2012). Subsequently, multiple alleles containing *gl_2_^arb^, gl_2_^b^, gl_3_^dav^*, *gl_3_^thur^, gl_3_^rai^, gl_3_^b^* (Bell and Stipanovic 1977), *Gl_2_^s^*(Barrow and Davis 1974),*Gl_2_^e^* (Kohel and Lee 1984; Zhang et al. 2001) and *Gl_2_^b^* (Zhu et al. 2004) associated with pigment gland formation were reported, the completely glandless character was controlled by two pairs of duplicate homozygous recessive genes (*gl_2_gl_2_gl_3_gl_3_*) or one dominant gene *GL_2_* . At the same time, Many transcription factors (TFs), such as *GhMYC2-like* (synonym *GoPGF* or *CGF3*), *CGF1*, *CGF2, RanBP2*, *GauGRAS1*, *GhNAC201* and *GhERF105*, were involved in the gland formation of cotton (Cai et al. 2003; Chang et al.2007; Cai et al. 2010; Cheng et al. 2016; Ma et al. 2016; Cai et al. 2019; Janga et al. 2019; Zhang et al.2020; Wu et al.2021). In the past few decades, some progress has been made in the molecular mechanism of gland formation. However,, little is known about the specific mechanism of pigment gland formation in cotton.

In this study, we described the properties and related functions of an ethylene response factor Named GhERF105a. The study provided a theoretical basis for analyzing the molecular mechanism of cotton gland formation, candidate gene and molecular-assisted selective markers for the breeding of new low-gossypol cotton varieties.

## Materials and Methods

### Plant materials and and treatments

CCRI12 (China Cotton Research Institute 12), CCRI17 (China Cotton Research Institute 17) and CCRI24 (China Cotton Research Institute 24) are Upland cotton cultivars with dark-colored pigment glands and high content of gossypol in both plants and seeds. While CCRI12XW, CCRI12YW, and CCRI17YW, which have glandless and low gossypol content in both seeds and plants, are glandless near isogenic lines (NILs) that differ nearly only in the gland trait of CCRI12 and CCRI17, respectively. The glandless trait of CCRI17YW and CCRI12YW are controlled by two the recessive genes *gl2* and *gl3*, while CCRI12XW is controlled by one dominant gene named *GL*_2_^e^. *GhMYC2-like*-silenced CCRI24 is a transgenic plant that silences *GhMYC2-like* gene. All materials were stored in Cotton Research Institute, the Chinese Academy of Agricultural Sciences (CAAS) (Anyang, China).

The seeds were immersed in water and followed by germination in a high humidity environment at 28°C in the dark for 2 d. Well-germinated seeds were subsequently planted in 0.3 litres pots of 7-cm diameter with one seed per pot in a commercially available sand/soil/fertilizer mix and grown for two to three weeks at 28°C (16 h light and 8 h dark) with LED lamps (Opple lighting Zhongshan China) in a greenhouse.

Samples from the different organs of both glanded and glandless cotton plants were collected and immediately frozen in liquid nitrogen and then stored at -80℃ for later use.

### Extraction of total RNAs

Total RNAs were isolated from all samples with liquid nitrogen using the RNAprep Plant RNA kit (polysaccharides&polyphenolics-rich) (TIANGEN BIOTECH (BEIJING)CO., LTD) according to the manufacturer’s instructions, The quantity and purity of RNAs were quantified photometrically using a NanoDrop OneC Microvolume UV-Vis Spectrophotometer with Wi-Fi (Thermo Fisher Scientific Inc., Waltham, MA, USA). Moreover, total RNA was electrophoresed using a 1.0% (w/v) denatured formaldehyde agarose gel to investigate its integrality.

### Synthesis of the first-strand cDNA

RNA was reversely transcribed into 1st strand cDNA in a 20 μL reaction volume using the PrimeSeript^TM^1stStrand cDNA Synthesis Kit (TaKaRa Bio, Dalian, China) following the manufacturer’s protocol of Reverse Transcription System. First, 2 μg of total RNA was mixed with 1.0 μL Oligo dT Primer (50 μm), 1.0 μL dNTP mixture (10mM each), then RNase free ddH_2_O was added to make the whole reaction volume up to 10μL, then, a total 10 μL reaction volume was incubated at 65℃ for 5 min and immediately placed on ice for 2 min to denature probable RNA secondary structure. Second, a mixture of 4 µL of 5xPrimeScript II Bufferr, 0.5 μL RNase Inhibitor(40 U/μL), 4.5 μL RNase free ddH_2_O and 1 μL Primescript 1I RTase (200 U/μL) were added to the chilled 10 µL reaction mixture prepared above and mixed. The mixture was incubated at 30℃ for 10 min, 42℃ for 60 min and terminated at 95℃ for 5 min.

### Identification and phylogenetic analysis of the GhERF105a gene in Gossypium spp

GhERF105a/d were identified by the comparative transcriptome analysis of the leaf of two pairs of glanded and glandless cotton accessions, which are L7 and L7XW, CCRI12 and CRI12XW. In order to study the character about *GhERF105a* gene in three cotton species, we downloaded the predicted protein sequences of *G. hirsutum* (Gh) (AADD), *G. barbadense* (Gb) (AADD), *G. arboreum* (Ga) (AA) and *G. raimondii* (Gr) (DD) from Cotton FGD (https://cottonfgd.org/about/download.html) and NCBI (https://blast.ncbi.nlm.nih.gov/Blast.cgi?PROGRAM=blastn&PAGE_TYPE=BlastSearch&LINK_LOC=blasthome). The protein sequences of *Arabidopsis thaliana* ERF were downloaded from TAIR database (https://www.arabidopsis.org/). The predicted molecular weight and isoelectric points of GhERF105 protein were calculated using the ExPASy program (http://web.expasy.org/protparam/). The phosphorylation sites were predicted using NetPhos3.1(http://www.cbs.dtu.dk/services/NetPhos/).

The glycosylation sites were predicted using NetNGlyc 1.0 (https://www.baidu.com/link?url=p4FgvTpgXkwkJk7d8K-wBXH8crmLp8QkywDjNmy98E33faOkACkzSct0fBq1lsn-Dh5UXqqbanc0eZhs_nI7OK&wd=&eqid=821afffb00016530000000025f2cc144). Multiple sequence alignments of the protein sequences were carried out using ClustalX (ver.1.81) with default settings, and Phylogenetic trees were constructed using MEGA 7.0 software by the Neighbour-Joining statistical method. The evolutionary distances were computed using the poisson correction method and the nodes of the trees were evaluated by boot-strap analysis with 1000 replicates.

### Molecular cloning of the full-length cDNA

The full-length cDNA of GhERF105 and other genes was amplified from the leaves of CCRI12 and cloned into the relative vector (such as pBI121, pGBKT7 et al.) as described previously (Wu et al 2021). The PCR products were electrophoresed on a 1% agarose gel, then were purified following the instructions in the QIAquick PCR Purification Kit (250) (Qiagen, Düsseldorf, Germany) and eluted in a final sample volume of 35 µL of Qiagen EB buffer. The primers used for cloning the cDNA of GhERF105a and GhERF105d were listed in Table S1.

### Gene Expression Analysis by Quantitative Real-Time PCR

Real-time quantitative (RT-qPCR) analysis was performed on the cDNA samples using the Premix Ex TaqTM (Tli RNaseH Plus) (TaKaRa Bio, Dalian, China) and ABI Quantstudio 5 Detection System (Applied Biosystems, Carlsbad, CA,USA). the reaction volume and conditions were described in previous study (Wu et al. 2021). The relative expression levels of all samples was normalized to that of the reference gene Actin (GenBank accession numbers: AY305733) and calculated according to the 2^−ΔΔCT^ method (Livak and Schmittgen 2001).The primers used for expression analysiswere listed in Table S1.

### Subcellular localization of GhERF105a

To study the subcellular localization of GhERF105a protein, the full length of GhERF105a without a termination codon was amplified and inserted into the *Xba*I and *Sma*Ⅰ sites of the binary vector pBI121-GFP to generate 35S-GhERF105a-GFP in-frame fusion with green fluorescent protein (GFP). The p35S-*GhERF105-GFP* plasmid and p35S-*GFP* (empty vector) were introduced into *Agrobacterium tumefaciens* strain GV3101 by the freeze-thaw method. The fusion construct or empty vector was introduced into the onion epidermal cells by *Agrobacterium*-mediated transformation. The onion epidermal cells had previously been incubated on MS agar plates in the light at 25℃ for 24 h. Transformed onion epidermal cells were cultured on the same MS media in the dark for 24 h at 25°C, then incubated at 25℃ for 48 -72 h in the light, followed by monitoring the localization of GFP with a confocal laser scanning microscope (Leica TCS SP8, Germany). The primers used for Subcellular localization of GhERF105a were listed in Table S1.

### Transactivation activity assay of GhERF105 protein

For yeast transactivation assay, the coding sequences (CDS) of *GhERF105a* and *GhERF105d* were cloned respectively into the EcoRI and NotI sites of pGBKT7 vectors to generate pGBKT7-*GhERF105a*/*GhERF105d* (Clontech, Palo Alto, CA, USA). The recombinant plasmid of pGBKT7-*GhERF105a/d* and plasmid with empty vector were transformed respectively into strain AH109 yeast cells according to the manufacturer’s instructions (Clontech, USA). The transactivation activity was determined by the growth on SD/-Trp and SD/-Trp/-X-a-gal (5-bromo-4-chloro-3-indolyl-α-D-galactopyranoside) plate. The images was taken after 96 h incubation at 30°C by stereomicroscope (Olympus SZX10, Japan). The primers used for transactivation activity were listed in Table S1.

### The promoter analysis of *GhERF105* genes

Total genomic DNA was extracted from young leaf tissue of the glanded cultivar CCRI12 using a modified CTAB (Cetyl Trimethyl Ammonium Bromide) method as described method (Paterson et al 1993), The 2-kb promoter region from the initiation codon of each *GhERF105a* and *GhERF105d* gene were amplified by PCR and inserted into the *Hind*Ⅲ and *BamH*I sites of the pBI121vector to generate pBI121- *GhERF105* construct. The promoter region of each *GhERF105a* and *GhERF105*d genes were analyzed by PlantCARE database (http://bioinformatics.psb.ugent.be/webtools/plantcare/html/) to reveal promoter cis-elements (Lescot et al 2002).The primers used for cloning the promoter were listed in Table S1.

### NMY51 assays

The full-length CDS of *GhERF105a* was amplified and cloned into the bait plasmid pDHBⅠ vector, and the full-length cDNA sequence of the gene (such as *Gh_A07G1044*) was cloned into the prey vector pPR3-N and co-transformed into yeast strain NMY51 cells together with pDHBⅠ-*GhERF105a* plasmid according to the Matchmaker user’s manual protocol (Clontech). Yeast transformants were selected by growth on synthetic dropout nutrient medium SD/-Leu/-His/-Ade/-Trp (Clontech) supplemented with 40mg/LX-α-gal (5-bromo-4-chloro-3-indolyl-a-D-galactopyranoside) at 30°C for 3 to 5 d to determine possible interactions between proteins. The images were taken by stereomicroscope (Olympus SZX10, Japan).The primers used for bait and prey recombinant plasmid construction were listed in Table S1.

### BiFC assay

For BiFC assays, the full-length CDSs of *GhERF105a* and *Gh_A07G1044* were amplified and cloned into pSPYNE and pSPYCE vectors, respectively, to generate N-terminal inframe fusions with nYFP and C-terminalin-frame fusions with cYFP constructs. The generated constructs of *GhERF105a* - nYFP and *Gh_A07G1044* - cYFP were transformed into the *A. tumefaciens* strain GV3101, and were then co-expressed in onion epidermal cells by *agrobacterium*-mediated transformation. After infiltration for 3 or 4 d, YFP fluorescence signals were detected by the fluorescence microscope (Olympus BX53, Japan). The primers used in BiFC experiment were listed in **Table S1.**

### Yeast one-hybrid assay

The coding sequence of *GhERF105a* was amplified and ligated into the *Sac*Ⅰ and *EcoR*I sites of the pGADT7 vector. The specific DNA fragments, GCC (AGCCGCCAGCCGCCAGCCGCC), mutated GCC (mGCC) (AGTTGCCAGTTGCCAGTTGCC) and DRE (TACCGACATTACCGACATTACCGACAT) were inserted into the pHIS2 vector. The plasmids were transformed into yeast strain Y187 and the transformants were selected on SD (-Trp/-Leu). Transformed colonies were subsequently grown on SD (-Trp/-His/Leu) medium with 80 mM 3-amino-1, 2, 4-triazole (3-AT), and were cultured at 30°C for 3 d. The primers used for the vector construction were presented in Table S1.

### Statistical analyses

The experiment was duplicated for three times, and the results were expressed as the mean values ± standard deviation (SD) of three biological and three technical replicates. Statistical significance of the data was evaluated using Student’s t-test (*P < 0.05, **P < 0.01).

## Results

### Cloning and sequence analysis

*GhERF105a* (Gene ID: *Gh_A12G1784*; Accession No. XP_016721164) and its homologous gene *GhERF105d* (Gene ID: *Gh_D12G1951*; Accession No. XP_016746305) were cloned from the leaves of CCRI12. The cDNA fragments of *GhERF105a* and *GhERF105d* were 711bp in length containing an open reading frame (orf) of containing initiation codon (ATG) and termination codon (TAA) (Fig. S1, Fig. 1). Sequence of amplified products for *GhERF105a* and *GhERF105d* was identical to the predicted database sequences respectively.The predicted proteins comprised of 236 amino acids containing a conserved GCC-box DNA-binding domain with the molecular mass of 26.3 kDa and theoretical isoelectric point of 7.72 and 7.57 (Table S2). the deduced protein of *GhERF105a* has a central AP2/ERF domain of 61 amino acids with two conserved amino acid residues, alanine (A) at position 14 and aspartic acid (D) at position 19 (Fig. S2), which is a typical characterization of the ERF-binding domain. However, 12 nucleotide mutation sites were discovered in between *GhERF105a* and *GhERF105d* gene, four of which were nonsynonymous mutation and 8 synonymous mutation (Fig. 1). The multiple sequence alignments showed that AP2/ERF domain was conserved between *GhERF105a* and other proteins (Fig. S3). The phylogenetic tree analysis indicated that GhERF105a protein shows a high similarity to GOBAR_AA13184 (100%, Accession No. KAB2053713), Ga_12G0841(99.6%, Accession No. XP_017636134), Gr008G214000 (97.9%, Accession No. XP_012439108) and GhERF105d (97.9%, Accession No. XP_016746305), respectively (Fig. 2a). The results from searching the Phyre2 database (http://www.sbg.bio.ic.ac.uk/~phyre2/html/page.cgi?id=index) showed a high similarity in the 3D structures between GhERF105a and GhERF105d (Fig. 2b). Sequence conservation suggested that the GhERF105a and GhERF105d belonged to class Ⅱ ERF subgroup of the AP2/ERF supfamily transcription factors.

**Fig. 1.**
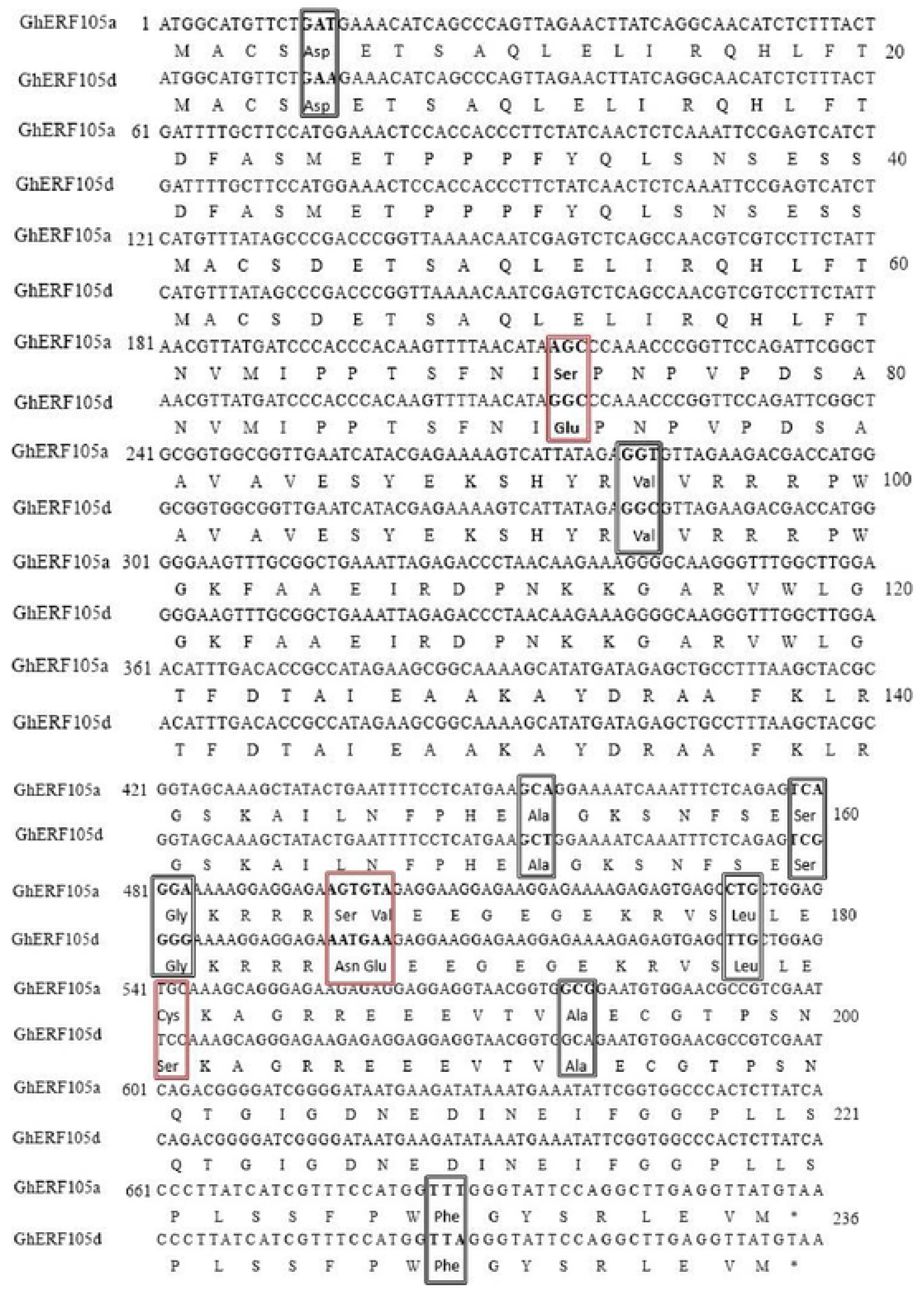
Length and its differences of nucleotide and amino acid between *Gh_A12G1784* (*GhERF105a*) and *Gh_D12G1951*(*GhERF105d*). Nucleotide positions are given at the left side of the sequence in the 5′ to 3′ orientation. The deduced amino acid sequence is shown beneath the nucleotide sequence and the amino acids are numbered to the right of the sequence. The nonsynonymous mutation and synonymous mutation are marked by red and black square frame respectively. The asterisk ‘*’ indicates the amino acid encoded by TAA.

**Fig. 2.**
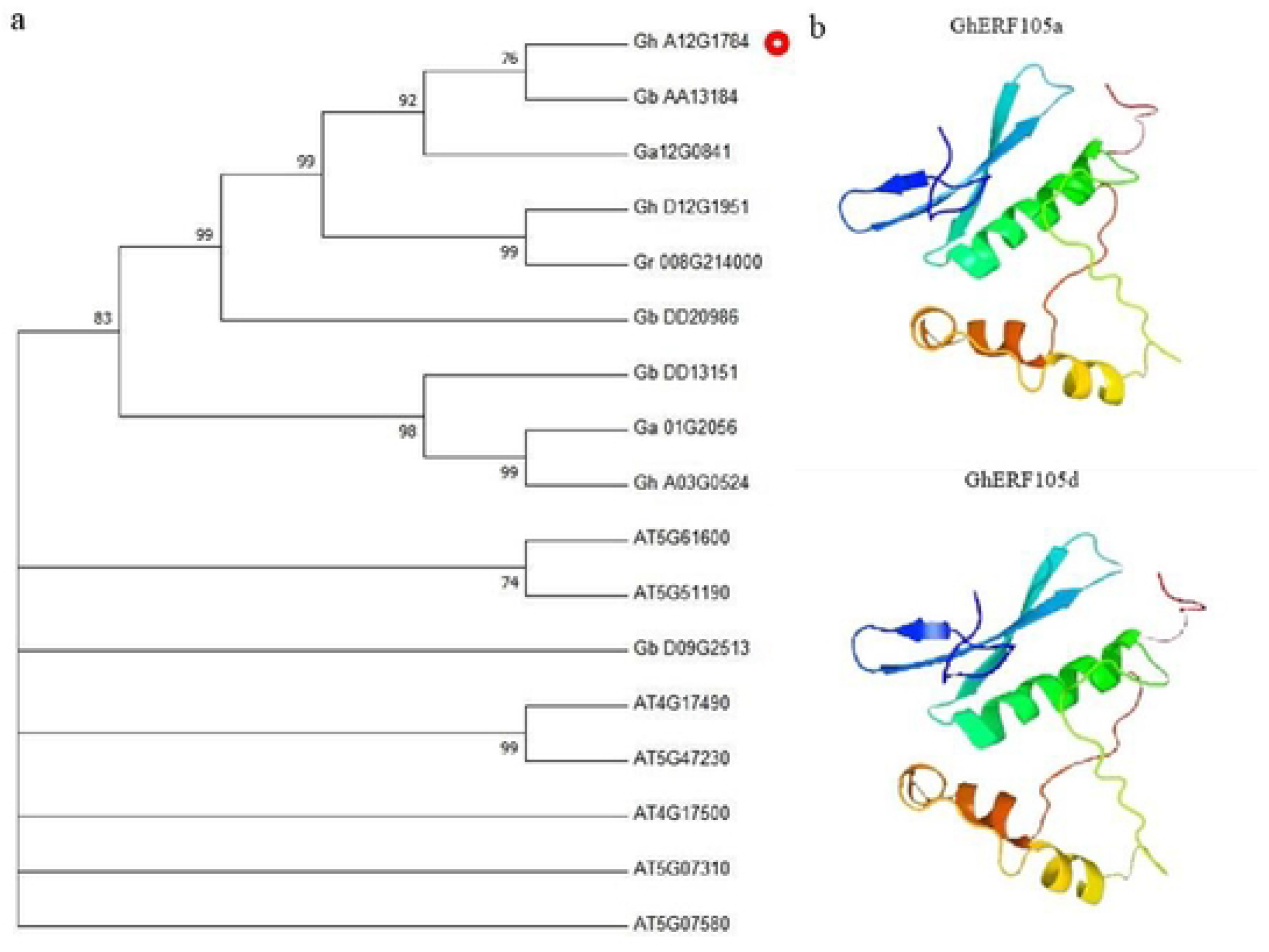
The phylogenetic tree and predicted three-dimensional structure of *GhERF105a* protein. (a) The phylogenetic tree was generated with the neighbor joining method using MEGA6.0 software. GhERF105a protein is indicated by the red circle. *Gh_A12G1784* (XP_016721164), *Gh_D12G1951* (XP_016746305), *Gh_A03G0524* (GhERF105), *Gb_D09G2513* (AAT77191), *Gb_AA13184* (KAB2053713), *Gb_DD13151* (PPD89903), *Gb_DD20986* (PPD82089), *Ga12G0841* (XP_017636134), *Ga01G2056* (XP_017640650), and *Gr008G214000* (XP_012439108) were derived from *Gossypium*, *AT4G17490*(NP_567529), *AT5G61600*(NP_200968), *AtERF105*(NP_568755.1), *AT5G47230*(NP_568679.1), *AT4G17500*(NP_567530.4), *AT5G07580*(NP_196375.3), *AT5G07310*(NP_196348.1) and *AT5G61590*(NP_200967.1) were derived from *Arabidopsis thaliana*. (b) Predicted three-dimensional structure of *GhERF105a* and *GhERF105d* protein. Black frame indicated identical amino acids between the *ERF* proteins.

Expression analysis of *GhERF105a/d*

Pigment glands are located on the surfaces of the stems, leaves, sepals, petals, and stigmas (McCarty et al. 1996), *GhERF105a* gene was associated with the development of cotton pigment gland (Wu et al. 2021). The results showed that transcription levels of *GhERF105a/d* are significantly higher in leaves and stems of glanded than those of glandless plants. At the same time, *GhERF105a* and its homologous gene *GhERF105d* showed the similar expression pattern in the different organs of glanded and glandless plants (Fig. 3,4).

**Fig. 3.**
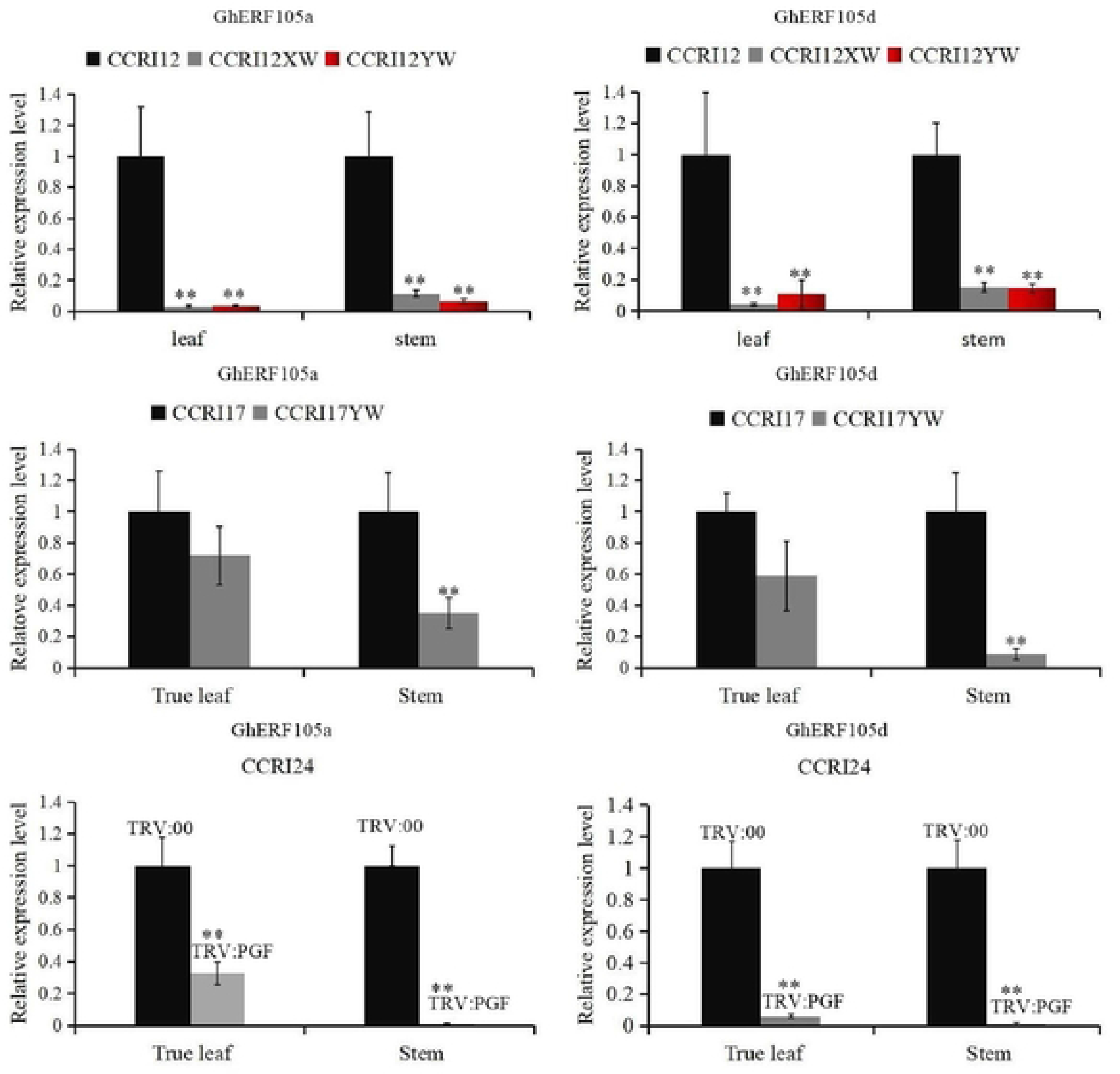
RT-qPCR expression analysis of GhERF105a gene and its homologous GhERF105d in the leaf and stem of CCRI12, CCRI12XW, CCRI12YW, CCRI17, CCRI17YW CCRI24, and *GhMYC2-like*-silenced CCRI24. The relative expression levels were calculated by the 2^−ΔΔCT^ method with the gene Actin (GenBank accession numbers: AY305733) as an internal control. The experiment was performed each with using three biological and technical replicates, and analyzed using Student’ t-tests (*P < 0.05, **P < 0.01). Error bars represent the standard of the mean values of three biological replicates. ‘*’ indicates the expression of *GhERF105* was significant difference in the leaf and stem of glanded plants and glandless plants.

**Fig. 4.**
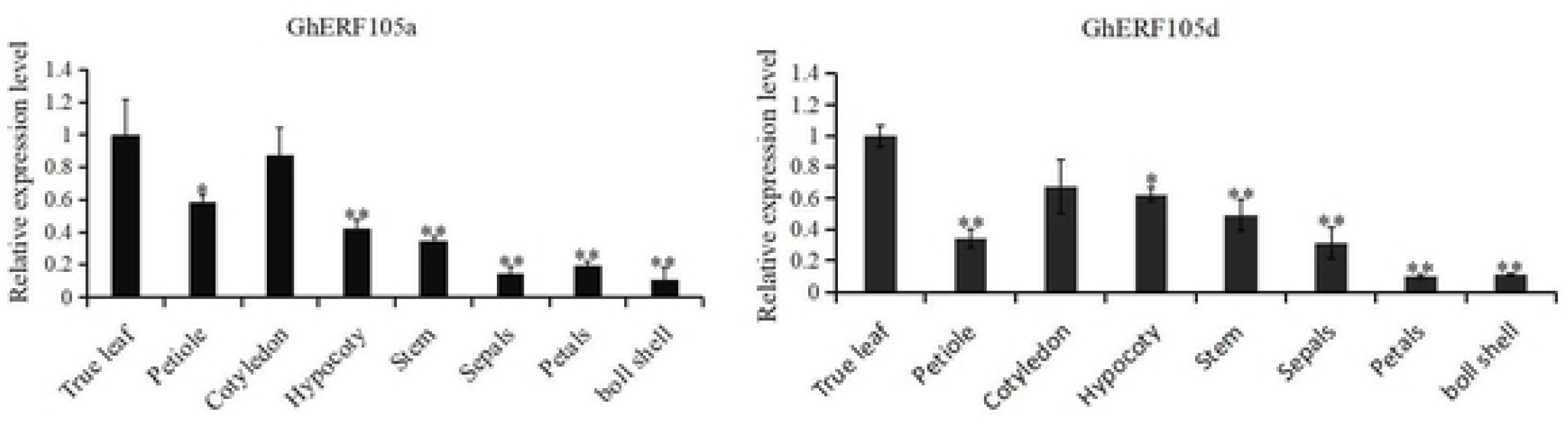
Spatiotemporal expression analysis of *GhERF105a* gene and its homologous *GhERF105d* in different organs of glanded plant. The relative expression levels were calculated by the 2^−ΔΔCT^ method with the gene Actin (GenBank accession numbers: AY305733) as an internal control. The experiment was performed each with using three biological and technical replicates, and analyzed using Student’ t-tests (*P < 0.05, **P < 0.01). Error bars represent the standard of the mean values (SE) of three biological replicates. symbol ‘*’ indicates the expression of GhERF105a/d were significant difference in the different organs of glanded cultivar.

### Subcellular localization of GhERF105a protein

As a transcription factor, one of the important features is its nuclear localization. The coding region of the *GhERF105a* gene was fused to the *GFP* gene and *GhERF105a-GFP* driven by the CaMV35S promoter was introduced into onion epidermis cells by *Agrobacterium*-mediated transformation. The fluorescence of GFP-GhERF105a was localized exclusively in the nucleus (Fig.5 b4-b6), whereas the fluorescence of GFP alone was observed in the whole cell (Fig.5 b1-b3). Therefore, the results confirmed that GhERF105a was a nuclear-localized protein.

**Fig. 5.**
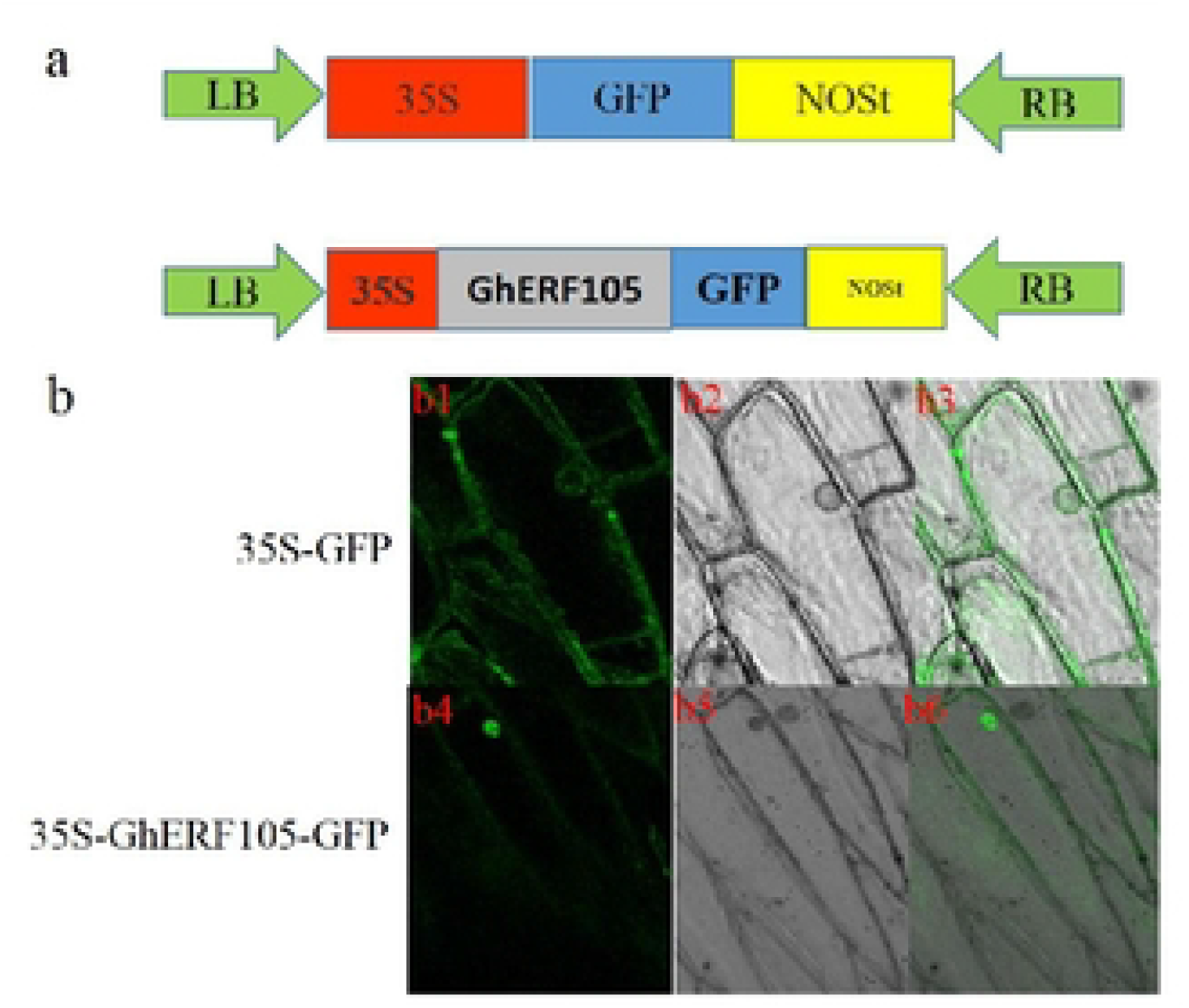
Subcellular localization of *GhERF105a* in cells. (a) Construction of *GhERF105-GFP* vector. (b) Subcellular localization of ***GhERF105a***-GFP. b1-b3, Transient transformation with 35S - GFP. b4-b6, Transient transformation with 35S-*GhERF105a*-GFP fusion construction. The photographs were taken in a dark field for green fluorescence (b1 and b4), in bright light for the morphology of the cells (b2 and b5) and in combination (b3 and b6). Bars 100 μm.

### Transactivation assay of GhERF105 proteins

To investigate the transcriptional activity of GhERF105a/d, *GhERF105a/d* cDNA were cloned into the EcoRI and NotI sites of pGBKT7 vector to generate *pGBKT7 - GhERF105* construct, the recombinant plasmid and the plasmid with an empty vector control were then transformed into yeast strain AH109 to analysis the transactivation activity respectively. The yeast strains transformed with the *pGBKT7-GhERF105a* and *pGBKT7-GhERF105d* were able to grow blue colonies on the selected medium SD/-Trp/-X-a-gal while those strains with empty vector pGBKT7 could grow white colonies (Fig.6). The result indicated that GhERF105a/d had the transcriptional activity.

**Fig. 6.**
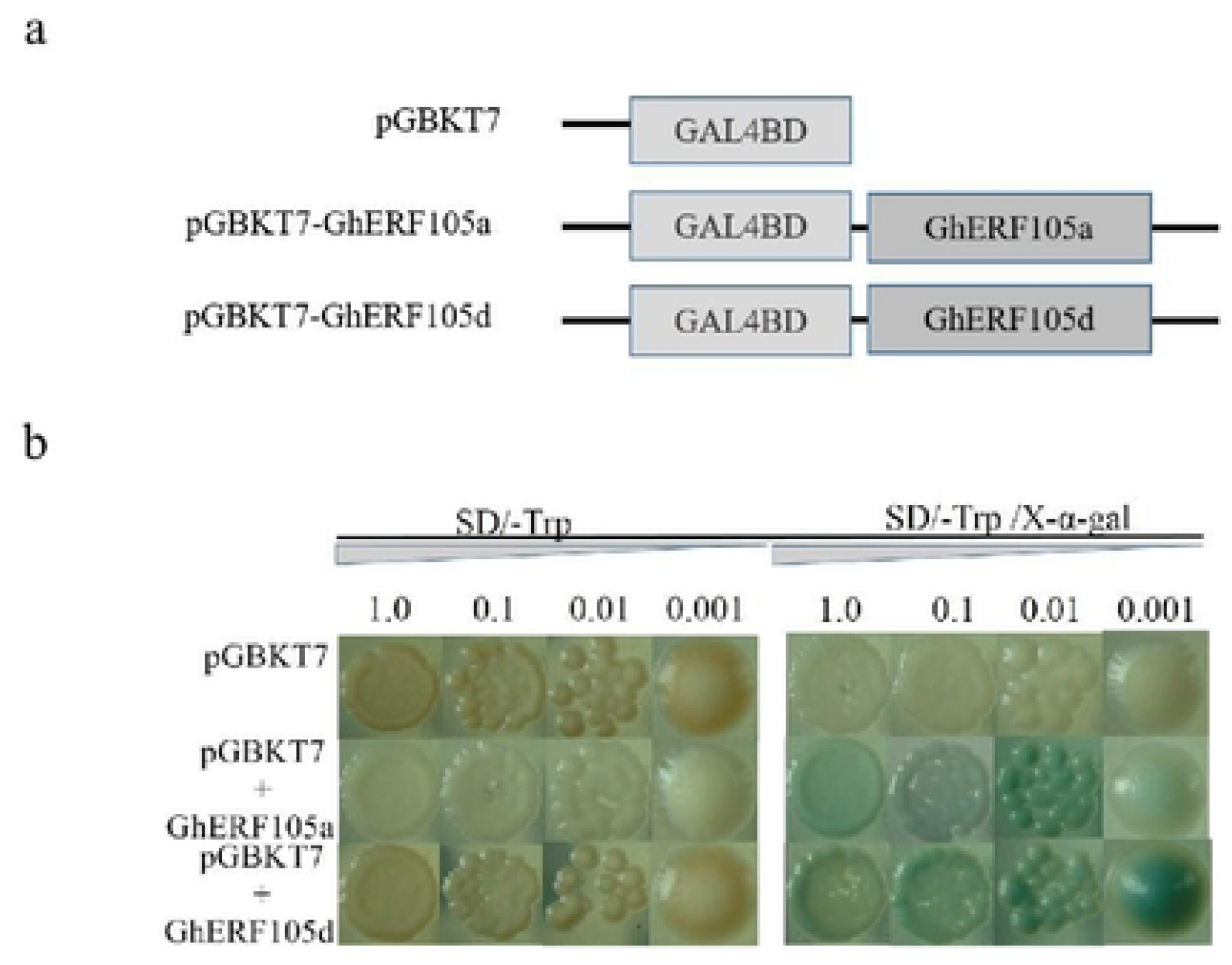
Transactivation activity of *GhERF105a* and *GhERF105d* in yeast cells. (a) Schematic diagram of the effector constructs used in the yeast assays. The effectors contained the GAL4 DNA binding domain coding region (GAL4BD) fused to GhERF105a, or GhERF105d. (b) the Yeast transformants with OD600 of 1.0, 0.1, 0.01 and 0.001 were selected by growth on the selected medium SD/-Trp/-X-a-gal at 30℃ for 4 d.

### Cis-element analysis of *GhERF105* genes in *G. hirsutum*

Promoter analysis is an effective method to study potential transcriptional regulation of genes. Therefore, To further investigate the functions of *GhERF105* genes, the 2 000-bp promoter fragment upstream of the *GhERF105a/d* initiation codon, as hypothetical promoter regions, were cloned and carried out using the PlantCARE (http://bioinformatics.psb.ugent.be/webtools/plantcare/html/) to reveal putative cis-elements. The result showed that there were many cis-elements in response to the development, phytohormone and stress in the promoters of GhERF105a/d (Fig.7). In addition, the eight most commonly cis-elements including the abscisic acid responsiveness element (ABRE), TC-rich repeats, G-box, GT1-motif, anaerobic induction element (ARE), heat induction element (STRE), TCCC-motif and WUN-motif were found in the promoters of GhERF105a/d (Fig.7). ARE, STRE, TC-rich repeats and WUN-motif are involved in defense and abiotic stress responsiveness (Kyozuka et al. 1994), while G-box, TCCC-motif and GT1-motif are involved in light response and development. In addition, ERE elements, which are involved in phytohormone responses, were found in the promoter region of GhERF105a. In brief, the above results suggest that GhERF105 genes may play important roles in the development, phytohormone response and stress response in cotton

**Fig. 7.**
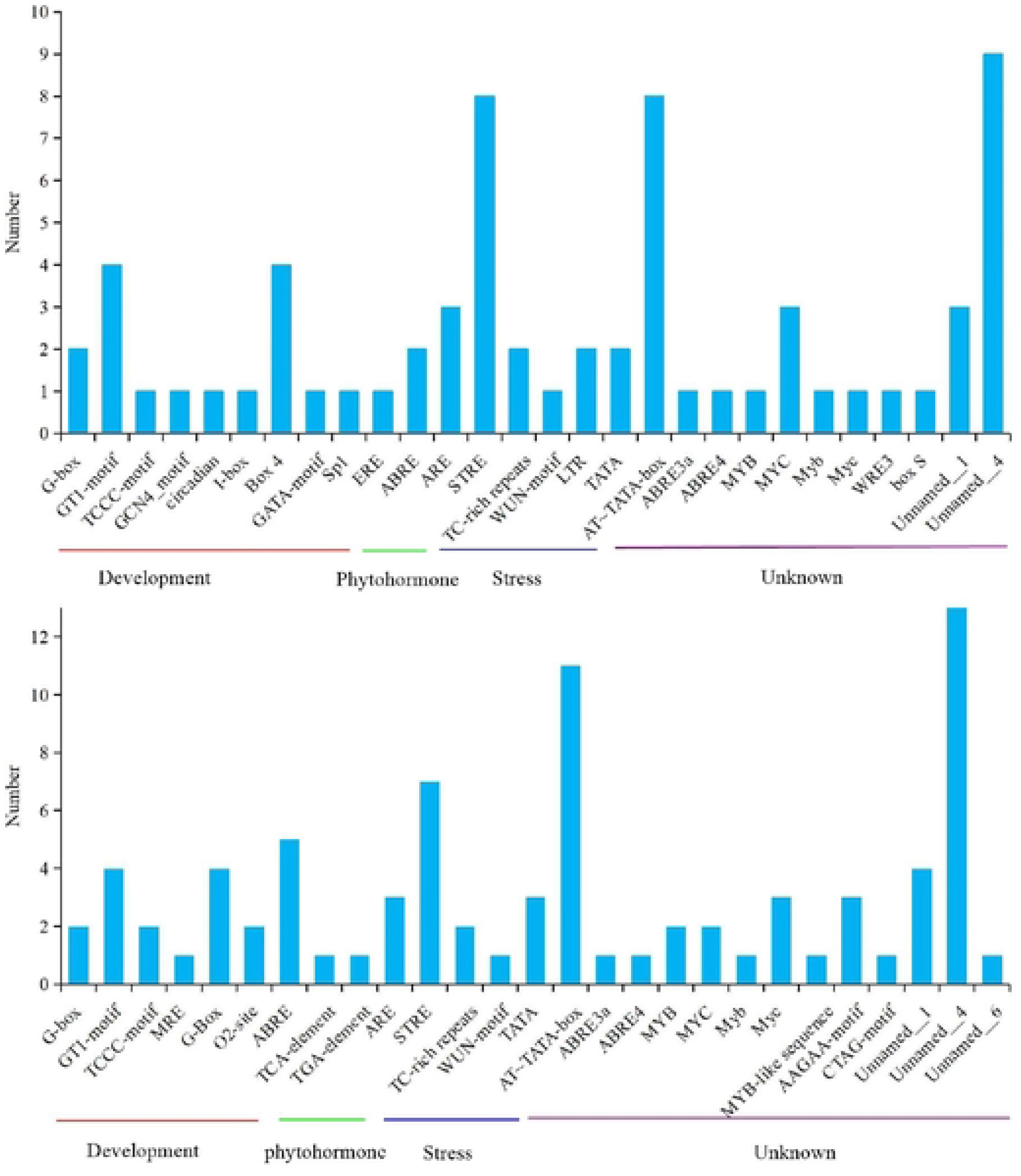
Cis-element analysis of *GhERF105*a and *GhERF105*d gene in *G. hirsutum*. The -2kb upstream sequence of the start codon of *GhERF105a/d* genes were analyzed by the PLANTCARE database.

### Interaction analysis between GhERF105a and Gh_A07G1044

Gh_A07G1044, as a putative protein interacted with GhERF105a, were screened from the cDNA library using the yeast two-hybrid system based on the split-ubiquitin. Therefore, to verify whether GhERF105a can interact with Gh_A07G1044 (Accession No: XM_016888374.1), *GhERF105a* was cloned in the vector pDHBⅠ (pDHBⅠ-GhERF105a), while *Gh_A07G1044* was cloned in the vector pPR3-N (pPR3-N-Gh_A07G1044). Yeast strains containing pDHBⅠ-GhERF105a and pPR3-N-Gh_A07G1044 were able to grow on the selective medium, while the yeast strains containing pDHBⅠ-GhERF105a and pPR3-N failed to grow (Fig. 8a). The results indicated that a specific interaction between GhERF105a and Gh_A07G1044.

**Fig. 8.**
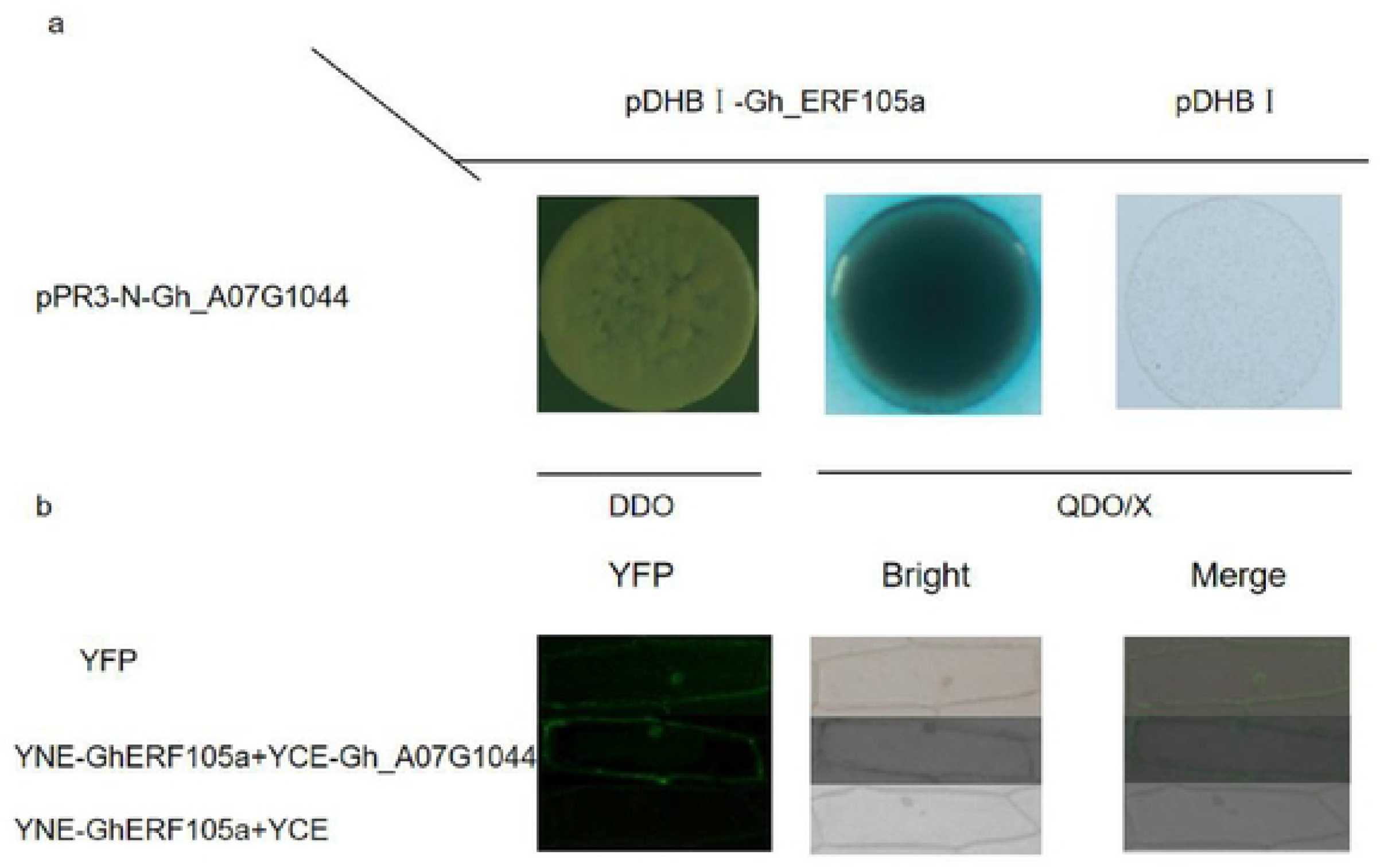
Protein interactions of *GhERF105a* gene in *G.hirsutum*. (a) Interaction patterns between *GhERF105a* and *Gh_A07G1044* proteins. The interactions were determined by growing on selective media SD/-Trp/-Leu (SD-2) and SD/-Trp/-Leu/-His/-Ade/X-a-Gal (SD-4). Bars = 200μm. (b) Test by BiFC for interaction between *GhERF105a* and *Gh_A07G1044* by fusing *GhERF105a* to the N terminus of YFP and *Gh_A07G1044* to the C terminus of YFP in onion epidermal cells. Bar = 200μm.

To further confirm that GhERF105a and Gh_A07G1044 can interact in plant, bimolecular fluorescence complementation (BiFC) was used as fluorescence would be revealed only if two proteins fused to split yellow fluorescent protein (YFP) interacted (Hu et al. 2002). For the BiFC assays, *GhERF105a* was fused to N-terminal YFP (*GhERF105a-nYFP*), while *Gh_A07G1044* was fused to C-terminal YFP (*Gh_A07G1044-cYFP*). The fluorescent signal was localized to the nuclear when *GhERF105a-nYFP* and *Gh_A07G1044-cYFP* were co-expressed in onion epidermal cells (Fig. 8b). The results suggested that GhERF105a indeed could interact in vivo with Gh_A07G1044 in plant.

### GhERF105a binds specifically to the GCC and DRE sequences

It was reported that ERF genes contain a conserved DNA-binding domain which can bind to GCC-box and DRE elements (Büttner and Singh 1997; Stockinger et al. 1997; Fujimoto et al. 2000; Zhang et al. 2004; Tanget al. 2007). To test whether GhERF105a specifically binds to GCC-box and DRE elements. As shown in Fig. 9, the bait strain (pHIS2-GCC) transfected with prey vector (pGADT7-GhERF105a) grew on SD (-Leu/-Trp/-His) medium containing 80 mM 3-AT, while mutant bait strain (pHIS2-mGCC) did not grow well. At the same time, the bait strain (pHIS2-DRE) transfected with prey vector (pGADT7- *GhERF105a*) could also grow on SD (-Leu/-Trp/-His) medium containing 80 mM 3-AT. These results showed that GhERF105a was able to interact with GCC-box and DRE elements.

**Fig. 9.**
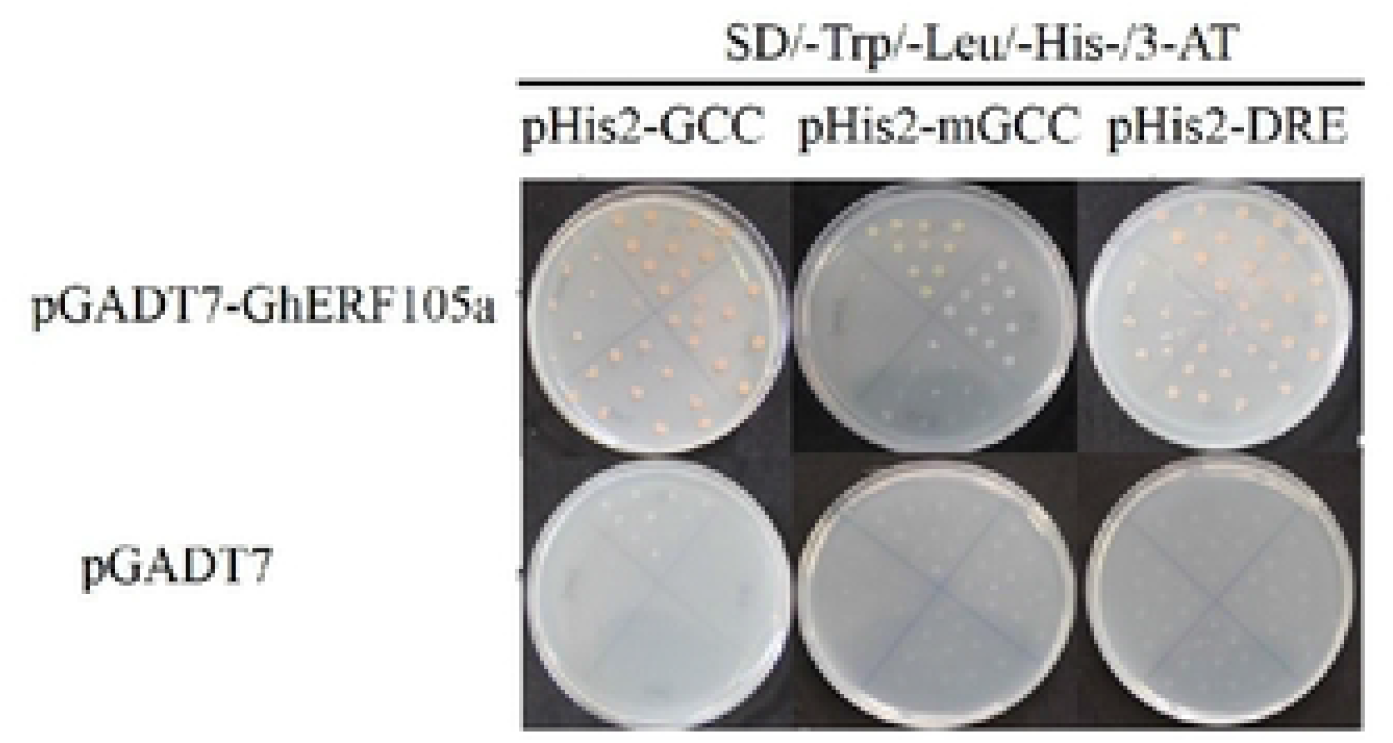
Binding to the GCC-box and DRE elements and transcription activity analysis of *GhERF105a*. The binding activity of *GhERF105a* to the GCC-box and DRE elements in yeast one-hybrid assay. Yeast cells were selected on SD (-Trp/-Leu/-His) media supplemented with 80 mM 3-AT.

## Discussion

Considering the importance of pigment glands for low-gossypol breeding of cotton, it is very significant to explore the molecular mechanism of the pigment gland formation. In the recent years, many genes related to pigment gland formation were identified and cloned from different cotton cultivars. *GhMYC2-like/GoPGF/CGF3* plays the most important role in pigment gland development (Cheng et al. 2016, Ma et al. 2016, Janga et al. 2019), *CGF1* shows similar functions to *CGF3*, and *CGF2* controls the density of pigment glands (Janga et al. 2019), the gossypol contents was decreased significantly in the new leaves of plants that were subjected to VIGS-mediated silencing of *CGF1*, *CGF2* or *GoPGF* (synonym *GhMYC2-like*, *CGF3*) genes (Ma et al. 2016, Janga et al. 2019). *GauGRAS1* controls the gland number of stems but does not change the form of glands in cotton leaves in *G. australe*, the content of gossypol in the stem of the *GauGRAS1*-silenced plants was significantly reduced (Cai et al. 2019). *GhERF105* controls the gland densiy of leaves but does not change the number of glands in the stems of cotton, The gossypol content in the leaves of the G*hERF105a*-silenced plants was significantly reduced (Wu et al. 2021). However, our understanding of the metabolic network related to gland formation and gossypol synthesis is still limited.

*ERFs* are among the most important TFs in the plant defense system (McCarty et al. 2005; Pre et al. 2008; Maruyama et al. 2013). In cotton, many *ERF* members have been cloned from cotton, such as *GhEREB2/3* (Duan et al. 2006), *GbERF2* (Zuo et al. 2007), *GhDREB1L* (Huang et al. 2007), etc. In this study, *GhERF105a* gene, which encoded a protein of 236 amino acids containing the AP2/ERF domain, was cloned from the leaves of CCRI12. Additionally, *GhERF105a* protein shares motif sites, such as different glycosylation and phosphorylation sites (Fig.S4; Fige.S5). These short regions might be required for transcriptional activation of *GhERF105a*. the phylogenetic tree analysis indicated that *GhERF105a* protein shows a high similarity to *GOBAR_AA13184* (100%, Accession No. KAB2053713), *Ga_12G0841*(99.6%, Accession No. XP_017636134), *Gorai_008G214000* (97.9%, Accession No. XP_012439108) and *GhERF105d* (97.9%, Accession No. XP_016746305).

To provide a greater insight into the biological roles of *GhERF105a* gene, the cis-acting elements in the promoter regions were analyzed with Plantcare datebase. Some potential regulatory elements associated with development, hormone and abiotic stress-related responses were present in the promoter regions of *GhERF105a* and its homologous gene *GhERF105d*, for example, G-box, TC-rich repeats and wun-motif was involved in light responsiveness, defense and stress responsiveness,and wound response,respectively. These elements may contribute to the corresponding functions. However, the analysis of promoter only provides some clues regarding the mechanism by which *GhERF105a* and *GhERF105d* genes respond to hormone transduction and stress signals. The function of these putative regulatory elements for the expression of *GhERF105a* and *GhERF105d* genes in cotton needs to be elucidated.

Expression pattern analysis can reveal the possible biological functions of target genes. RT-qPCR analysis has demonstrated that the transcription levels of GhERF105a/d are significantly higher in leaves and stems of glanded than those of glandless plants. The expression of *GhERF105a/d* were higher in CCRI24 than that *GhMYC2-like*-silenced CCRI24. In addition, GhERF105a/d were expressed constitutively in different organs (e.g., cotyledon, hypocotyl, petiole, leaf, stem, sepals, petals and boll shell) but were abundantly expressed only in leaves of cotton. RT-qPCR analysis indicated the *GhERF105a* and its homologous gene *GhERF105d* showed similar expression patterns in the leaves and stems of glanded and glandless plants. The results indicated that *GhERF105a/d* might be involved in gland formation.

*ERF* genes have been shown to act as transcription factors and are able to bind to GCC-box and (or) DRE elements in previous studies (Büttner and Singh 1997; Stockinger et al. 1997; Fujimoto et al. 2000; Tournier et al. 2003; Lee et al. 2004; Zhang et al. 2004; Tang et al. 2007). such as *ORA59* (Zarei et al. 2011), *AtERF6* (Wang et al. 2013), *AtERF96* (Catinot et al. 2015, *StERF3* (Tian et al. 2015), *DREB1A/2A*(Stockinger et al. 1997) and *DREB1A/CBF3 and DREB2A* (Sakuma et al. 2002). In this study, *GhERF105a* could bind specifically to the GCC-box and DRE elements and showed the transactivation activity (Fig.9). In combination with its nuclear localization, the result indicated that *GhERF105a* may function as a transcription activator and regulate defense-related genes expression by binding to GCC-box elements present in their promoters.

YABBY2 transcription factor regulate leaf development and are involved in stress response of the plant. It was reported that *ATCRC,*one of the YABBY2 transcription factors, is expressed in the distal end of the nectaries, and it mainly regulates the development of carpels and nectaries in *Arabidopsis* (Eckardt 2010; Zhang et al. 2013; Ha et al. 2010). Silencing *GaNEC1* changed obviously the morphology of nectarine cells in cotton varieties, the result demonstrated that the gene was involved in regulating the development of nectariesa and indirectly defended against pests in cotton (Hu et al. 2020). The yeast split-ubiquitin assays and BiFC confirmed that *Gh_A07G1044*, which is a member of YABBY2 transcription factors, could interact with GhERF105a in vivo. It was speculated that the interaction may be involved in the process of cotton pest control of leaves. These findings need to be confirmed by further experiments.

In addition, there are minor changes in predicted protein molecule, protein isoelectric point and the type and number of promoters, whereas the gene length and expression pattern are highly similar between *GhERF105a* and *GhERF105d* (Table S2, Fig.3, 4). In particular, *GhERF105a* shares similar structure and protein properties with *GhERF105d*, with a conserved AP2 domain. This result demonstrates that homologous genes originating from the progenitors can evolve independently at the same rate with few changes.

In conclusion, the cloning and characterization of *GhERF105a* both provide new information to study the molecular mechanism of gland formation and other biological functions in upland cotton.

## Conclusion

Ethylene-responsive factors (ERFs) are important regulators of plant gene expression. In this study, *GhERF105a/d* were cloned from cotton cultivar CCRI12. RT-qPCR analysis demonstrated *GhERF105a/d* were expressed at higher levels in the leaves and stems of glanded plants, and the transcription levels of *GhERF105a/d* were significantly higher in the leaves of CCRI24 plants than that of *GhMYC2-like*-silenced CCRI24. In addition, GhERF105a is constitutively expressed in different organs but were abundantly expressed only in leaves of cotton. Transient expression analysis using GhERF105-GFP demonstrates that GhERF105 is specifically expressed in the nucleus of onion epidermal cells and could bind specifically to GCC box and DRE element in yeast one-hybrid system. Promoter analysis also indicated that the 2000-bp upstream region of *GhERF105a* possessed some elements associated with development, hormone and abiotic stress-related responses. Collectively, these results indicated that GhERF105a might be involved in the gland formation and functioned as positive trans-acting factors in the plant responses to environmental stress.

## Author contributions statement

C.F.W conceived the research, performed the experiments and draft the manuscript; S.Y.L helped to prepared figures, provided the discussion and propose. X.M.M supervised all of the project and corrected the manuscript. All authors read and approved the final manuscript.

## Acknowledgements

We are grateful to the State Key Laboratory of Cotton Biology, Institute of Cotton research, Chinese Academy of Agricultural Sciences for providing the seeds of *G. hirsutum*, and the base vectors for assembling the transformation constructs. This research is supported by the PhD Start-up Fund of Natural Science Foundation of Anyang Institute of Technology (No. BSJ2022034).

## Abbreviations

BiFC: Bimolecular Fluorescent Complimentary
ERF: Ethylene transcription factors
GFP: Green Fluorescent Protein

## Availability of data

All raw sequencing data in the study have been deposited in the Short Read Archive (SRA) (www.ncbi.nlm.nih.gov/sra). The accession numbers are SRR1652340, SRR1652393, SRR1652399 and SRR1652403 available at https://www.ncbi.nlm.nih.gov/sra/. In addition,the accession number of genes in the study were listed in Table. S3

## Supplementary figures

**Fig. S1. Amplification of the full-length cDNA of *GhERF105s***. lane 1: DNA marker(MarkⅢ); lane 2 and 3: the full-length cDNA of *GhERF105a* and *GhERF105d*

**Fig. S2. The deduced amino acid sequence of Upland cotton GhERF105a protein**. The conserved ERF domain is underlined and marked in black bold. The asterisk ‘*’indicates the amino acid encoded by TGA; The alanine and aspartic acid residues at positions 14 and 19 in the AP2/ERF domain are marked by red triangles.

**Fig. S3. The multiple sequence alignment of GhERF105 with other ERF proteins from Arabidopsis**. The sequence alignment was performed using ClustalX (1.81) software. The AP2 DNA binding domain is indicated by purple box. Dashes show gaps in the amino acid sequences introduced to optimize alignment. *Gh_A12G1784* (XP_016721164), *Gh_D12G1951* (XP_016746305), *Gh_A03G0524* (GhERF105), *Gb_D09G2513* (AAT77191), *Gb_AA13184* (KAB2053713), *Gb_DD13151* (PPD89903), *Gb_DD20986* (PPD82089), *Ga12G0841* (XP_017636134), *Ga01G2056* (XP_017640650), and *Gr008G214000* (XP_012439108) are derived from *Gossypium*, *AT4G17490*(NP_567529), *AT5G61600* (NP_200968), *AtERF105*(NP_568755.1), *AT5G47230* (NP_568679.1), *AT4G17500* (NP_567530.4), *AT5G07580* (NP_196375.3), *AT5G07310* (NP_196348.1) and *AT5G61590* (NP_200967.1) are derived from *Arabidopsis thaliana*, The square frame represents conserved domain.

**Fig. S4. The predicted result of potential phosphorylation of *GhERF105a* protein**. GhERF105a protein contains twenty-three serine (S) phosphorylation sites, two tyrosine (Y) phosphorylation sites, and seven threonine (T) phosphorylation sites.

**Fig. S5. The predicted result of potential glycosylation of GhERF105a protein**. *GhERF105a* protein contains one glycosylation site at position 156.

**Table S1** List of primers used in the experiment

**Table S2** Comparison of physicochemical properties of GhERF105a and GhERF105d genes

**Table S3** List of the name and its accession number of each gene used in the study

